# Varmole: A biologically drop-connect deep neural network model for prioritizing disease risk variants and genes

**DOI:** 10.1101/2020.03.02.973743

**Authors:** Nam D. Nguyen, Daifeng Wang

## Abstract

Population studies such as GWAS have identified a variety of genomic variants associated with human diseases. To further understand potential mechanisms of disease variants, recent statistical methods associate functional omic data (e.g., gene expression) with genotype and phenotype and link variants to individual genes. However, how to interpret molecular mechanisms from such associations, especially across omics is still challenging. To address this, we develop an interpretable deep learning method, Varmole to simultaneously reveal genomic functions and mechanisms while predicting phenotype from genotype. In particular, Varmole embeds multi-omic networks into a deep neural network architecture and prioritizes variants, genes and regulatory linkages via drop-connect without needing prior feature selections. Varmole is available as a Python package on GitHub at https://github.com/daifengwanglab/Varmole.

## Introduction

Statistical analyses have associated a variety of genomic variants with phenotypes in human diseases such as cancers and brain disorders. For example, genome-wide association study (GWAS) analyses have identified a number of disease risk single-nucleotide polymorphisms (SNPs) from population genetic data. Furthermore, to determine the respective contribution of individual associated variants to disease phenotypes, polygenic risk scores (PRSs) have been developed based on linear regression model to quantify how much proportions of phenotypes can be explained by specific variants. However, like other complex polygenic diseases and possibly under an omnigenic model [1], hundreds even thousands of variants and genes are expected to function together as networks in diseases, with each individual risk gene contributing small effect. Therefore, interpreting such statistical scores are still challenging for discovering causal variants (e.g., a majority of GWAS SNPs are on noncoding regions) and understanding underlying mechanisms of risk variants.

To address this, functional information from omics has been integrated to fine-map and link variants to molecular and cellular activities, aiming to reveal causal variants and mechanisms. For example, a number of molecular quantitative trait loci (QTLs) have associated variants with molecular activities (e.g., eQTLs for gene expression). Also, the imputation methods such as TWAS [2] and PrediXcan [3] use these QTLs to impute molecular activities from disease variants and then connect to corresponding disease phenotypes. However, such associating variants, molecular activities and phenotypes at separate steps, each of which likely use different population data, potentially mismatch molecular activities and disease phenotypes. For example, only between 3.5% and 11.7% of top eQTLs in the Genotype-Tissue Expression (GTEx) project are the actual causal variants affecting gene expression across various human tissues [4]. Using GTEx eQTLs to impute gene expression for brain diseases may not be able to discover disease-specific expression and regulation. Thus, to discover underlying causal mechanisms, prior biological knowledge has been embedded into machine learning models for genotype-phenotype prediction [5, 6].

Therefore, we develop an interpretable deep learning method, Varmole to (1) predict disease phenotypes from genotype and molecular activities in a single, coherent model; (2) simultaneously prioritize causal variants and molecular mechanisms such as gene regulation for specific phenotypes. To achieve these, Varmole embeds prior biological knowledge such as QTLs and gene regulatory networks into a deep neural network; i.e. defining biological architecture so that the model is interpretable, rather than conventional fully connected “black box” neural network. Furthermore, compared to our previous work that requires selecting genes and variants before training and binarized gene expression data [5], Varmole is more scalable for enabling the implicit feature selection via Lasso regularization and drop-connect and as well as taking input data with continuous values.

## Methods and Implementation

Varmole is built on a deep neural network model with multiple hierarchical layers corresponding to particular biological or clinical types. It aims to predict phenotypes from genotypes indirectly mediated by gene expression. Unlike conventional neural networks, Varmole (Figure 1A) is a grey box model in which partial internal structure is defined by prior biological knowledge; i.e., a biologically interpretable model. The first layer of Varmole is a transparent layer of genes and genetic variants (e.g., SNPs) modeling the relationships between genotype and gene expression such as eQTLs [5]. The first hidden layer models genes. We then use the gene regulatory network (GRN) to build the connectivities between the first layer and the first hidden layer, modeling gene regulatory relationships among genes such as transcription factors (TFs) to target genes. That said, the extra GRN structure will be added to the transparent units of Varmole to reflect regulatory relationships between various genomic elements to help uncover the molecular mechanisms between genotype and phenotype. The transparent layer of Varmole enables the interpretation in which important features (i.e., SNPs, genes) and important paths (i.e., SNP-gene and gene-gene regulation relationships) contributing to the prediction results are extracted. The other layers of Varmole model the mechanisms in which gene expression gives rise to the phenotypes. Furthermore, Varmole does not have to input prior select features.

**Figure 1.**
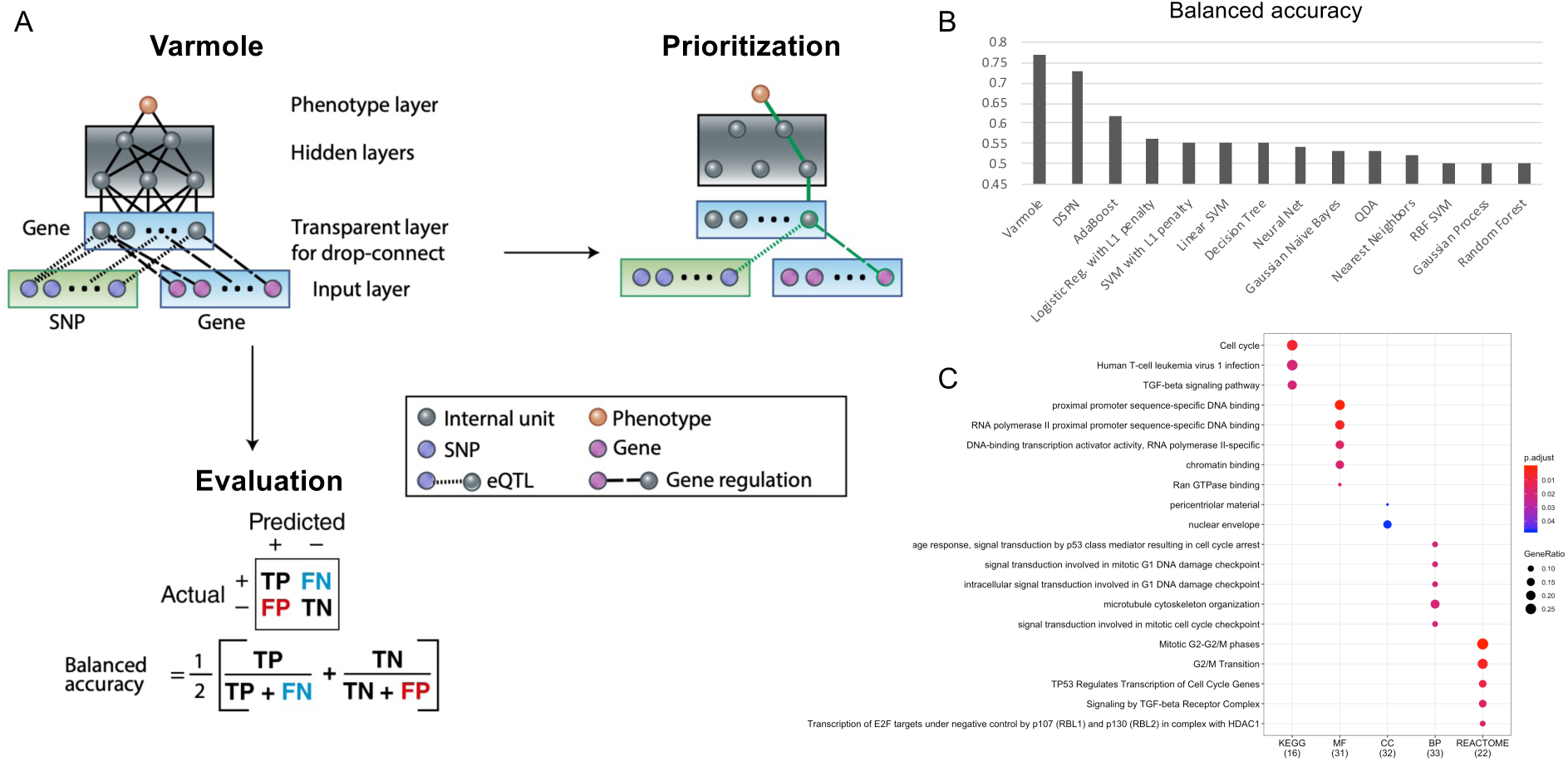
Varmole, a biologically drop-connect-based deep neural network model for prioritizing disease risk variants and genes. (A) Varmole model has four layers: Layer 1 inputs SNPs and genes; Layer 2 is a transparent layer duplicating the gene nodes in Layer 1; Layer 3 has multiple hidden layers; Layer 4 is the phenotype layer. The prior biological networks from eQTLs and GRNs link SNPs/genes to genes from Layer 1 to Layer 2, which enables biologically drop-connect. Varmole can be evaluated by balanced accuracy and prioritizes the genes, SNPs and links for a phenotype (e.g., green path) using the integrated gradients. (B) Classification performance comparison by the balanced accuracy between Varmole and other state-of-the-art methods for genotype-phenotype prediction including our previous DSPN model [5]. (C) Enrichment of top prioritized genes for predicting schizophrenia by clusterProfiler [15].

Instead, it uses the DropConnect [7], a *ℓ*-1 LASSO-based regularization technique generalized from DropOut architecture for prioritizing SNPs, genes and connections while predicting phenotypes; i.e., implicit feature selection. DropOut and DropConnect are the two most effective regularization techniques specifically for deep learning models which are based on a random subset selection of output activations (in case of DropOut) or of weight connections (in case of DropConnect) between the two consecutive layers of a neural network. Instead of selecting a subset of connections randomly as conventional DropConnect, the connections in Varmole are selected according to prior biological knowledge (i.e., GRNs and eQTLs); i.e., Biology-based DropConnect. In addition to interpretability, this architecture also helps address the overfitting issues due to the problem of a small number of samples vs. a large number of features in the biological datasets. Particularly, Varmole is composed of the following components:

1. Input *X* is the concatenation of gene expression matrix *E*^*n×p*^ (*n* genes by *p* samples) and SNP genotype (e.g., dosage) matrix *G*^*m×p*^(*m* SNPs by *p* samples) *X*^(*m*+*n*) *×p*^ *=* [*G*^*T*^*E*^*T*^]^*T*^, where (.)^*T*^ is transpose;
2. The first transparent layer *Z*_1_ with its dimension being the number of genes *n*. Hence the weight matrix *W*_1_ of this layer has a dimension of (*m+n*) *× n*. The links of eQTLs and GRNs are also embedded in this layer. Specifically, the two binary matrices *A*^*m×n*^ and *B*^*n×n*^ encode eQTL and GRN respectively where *A*_*i,j*_=1 if SNP *i* is associated with Gene *j* in eQTL data, and *B*_*i,j*_=1 if Gene *i* regulates Gene *j* in gene regulatory network. Instead of a fully connection between input *X* and hidden layer *Z*_1_, we now have a “Biology-based DropConnect” layer *Z*_1_ *= f*(*X*^*T*^(*W*_*1*_ ⊙ *C*)*+b*_*1*_), where *C =* [*A*^*T*^*B*^*T*^]^*T*^, where ⊙ is Hadamard product. Varmole enables an implicit feature selection method while training with *ℓ*-1 penalty (Lasso) to *W*_*1*_;
3. Other fully connected hidden layers *Z*_*l*_ indexed by *l ∊ 2* … *L* – *1*;
4. Softmax classification layer 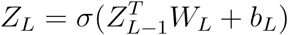;
5. The Cross Entropy Loss is used to quantify the classification error:

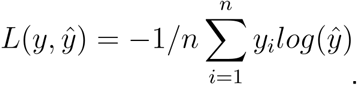

Once the Varmole model is trained, we can further use a derivative-based method called integrated gradient [8] for prioritizing nodes (i.e., SNPs and genes) and connections (i.e., eQTLs and regulatory relationships) for phenotypes. In particular, we computed the gradient of the model’s prediction with respect to each individual SNP and/or gene to show how the output response value (i.e., disease vs. control) changes with respect to a small change in input gene expression and/or SNP genotype. Hence, calculating these gradients for given input SNPs and genes provides potential clues about which SNPs and/or genes attributed the disease outcomes. This can be also interpreted to see which features are not selected due to *ℓ*-1 regularization since the gradients for these input features are zeros. Furthermore, the gradient for given input features are also decomposed (via the chain rule) into a flow of gradients’ attribution via a visible unit encoding the expression of a gene in the second layer for prioritizing the importance of paths from SNPs to genes (i.e., eQTLs) or from genes to genes (i.e., regulatory relationships). For example, a particular Varmole model can input to the Python package, Captum [9] that implements the integrated gradient method for prioritizing SNPs, genes and links.

## Usage case

We applied Varmole to the functional genomic data for the human brain in PsychENCODE consortium [5] for predicting schizophrenia from SNP genotype dosage and gene expression at the population level. After filtering out SNPs and genes to match with those in eQTLs and GRN, we are left with 771667 SNPs and 10450 genes. We further filter out those genes and SNPs such that each SNP in eQTLs has at least four associated genes. Finally, we input the genotype data of 27472 SNPs (i.e., dosage) and the expression data of 1493 genes from 487 schizophrenia and 891 non-schizophrenia samples. Also, we used a full GRN in PsychENCODE for the human brain to connect genes in Varmole. We also used 97210 eQTLs in PsychENCODE to link 27472 SNPs and 1493 genes in Varmole. The input data was splitted into train/validation/test with the ratio 64/16/20. After hyperparameter tuning, we came up with a deep neural network with ReLU activation functions consisting of: (1) the input layer containing 28965 features (1493 genes + 27472 SNPs); (2) the layers above containing 1493×27472, 1000×1493, 500×1000 and 500×1 connections, respectively; (3) the output layer predicting schizophrenia by a probability (i.e., via a single-unit with a sigmoid activation). Other hyperparameters were optimized as follows: Train batch size = 50; Learning rate = 0.001 and Weight decay = 0.01 (with Adam method [10]); *ℓ*-1 penalty parameter: 0.0001; Number of training epoch: 60.

While the conventional classification accuracy is the typical metric for measuring the performance of a classification algorithm, it potentially misleads with respect to data imbalance since an imbalanced training set may bring about a classifier that is biased towards the more frequent class. For example, in binary classification, the classifier can label every test point to the dominant class and thus yield an optimistic accuracy estimate. To address this, the balanced accuracy (BACC) [11] has been developed to address imbalanced training data by taking the average of sensitivity (true positive rate) and specificity (true negative rate). In particular, BACC balances the contributes of different classes by assigning a weight to each class such that the resulted classifier can learn equally from all classes (Figure 1A). The BACC also equals the area under the ROC curve with binary predictions. Also, if the dataset is balanced, this metric is equivalent to the regular accuracy. Therefore, given imbalanced sample sizes across disease and control in this application, we used the balanced accuracy to evaluate Varmole and found that it outperforms other state-of-the-art classifiers (BACC=0.77, Figure 1B). In addition, we further used the integrated gradient based method, Captum to prioritize the genes for predicting schizophrenia, based on the neural network architecture. In total, we ranked the importance of 27472 SNPs, 1493 genes, 97210 SNP-gene, and 3446 gene-gene regulatory links for predicting schizophrenia from their Captum scores (Supplemental Tables 1 and 2). We found that top prioritized genes (e.g., N=43 with Captum score > 0.01) are enriched with gene regulation, DNA damage and cell cycle functions (Figure 1C), suggesting them as potential biomarkers for schizophrenia. This result was also supported by experimental studies [12-14].

## Conclusion

This paper introduces an interpretable deep learning tool, Varmole to predict disease phenotypes from multi-omic data. Using embedded biological networks for drop-connect, Varmole simultaneously prioritizes genomic variants and functional mechanisms such as gene regulation for particular phenotypes. This provides the biological interpretability of Varmole, compared to that many existing machine learning methods attempt to learn a “black box” between genotype and phenotype. Varmole is publicly available as a Python package at Github, https://github.com/daifengwanglab/Varmole.

